# AE-LGBM: Sequence-Based Novel Approach To Detect Interacting Protein Pairs *via* Ensemble of Autoencoder and LightGBM

**DOI:** 10.1101/2020.07.03.186866

**Authors:** Abhibhav Sharma, Buddha Singh

## Abstract

Protein-protein interactions (PPIs) are a vital phenomenon for every biological process. Prediction of PPI can be very helpful in the probing of protein functions which can further help in the development of new and powerful therapy designs for disease prevention. A lot of experimental studies have been done previously to study PPIs. However, lab-based experimental studies of PPI prediction are resource-extensive and time-consuming. In recent years, several high throughput, computational approaches to predict PPI have been developed but they could be fallible in terms of accuracy and false-positive rate. To overcome these shortcomings, we propose a novel approach AE-LGBM to predict the PPI more accurately. This method is based on the LightGBM classifier and utilizes the Autoencoder, which is an artificial neural network, to efficiently produce lower-dimensional, discriminative, and noise-free features. We incorporate conjoint triad (CT) features along with Composition-Transition-Distribution (CTD) features into the model and obtained promising results. The ten-fold cross-validation results indicate that the prediction accuracies obtained for Human and Yeast datasets are 98.7% and 95.4% respectively. This method was further evaluated on other datasets and has achieved excellent accuracies of 100%, 100%, 99.9%, 99.2% on E.coli, M.musculus, C.elegans, and H.sapiens respectively. We also executed AE-LGBM over three important PPI networks namely, single-core network (CD9), the multiple-core network (The Ras/Raf/MEK/ERK pathway), and the cross-connection network (Wnt Network). The method was successful in predicting the pathway with an impressive accuracy of 100%, 100%, and 98.9% respectively. These figures are significantly higher than previous methods that are based on state-of-the-art models and models including LightGBM or Autoencoder, proving AE-LGBM to be highly versatile, efficient, and robust.

## 2. Introduction

Protein-Protein Interaction (PPI) has always possessed an important status in the scientific domains of proteomics due to its key role behind the underpinning of every biological process. The predictions of protein-protein interactions were initially performed only through stately lab-based experimental techniques expanding from affinity chromatography to co-immunoprecipitation [1]. Methods such as yeast two-hybrid screening [2-3], X-ray crystallography, protein chips [4], and affinity purification [5] have been used to study PPI at a molecular level and have generated a substantial amount of data of potentially interacting protein pairs. However, these lab-based experimental techniques remained labor-intensive and time-consuming processes and have fallible performance scores. To overcome these inadequacies many computational methods have been developed for the prediction of PPIs based on the structural, biological, and genomic properties of biomolecules[6-8]. Probing into these proposed methods, it is very convincing that over the time, machine learning-based models have performed significantly better in terms of the time expanse, accuracy, and the false prediction rate than that of the wet-lab based PPI analysis [9-11]. W. Chen et al [12] proposed a method based on a random forest classifier. The works of Shen et al [13] developed a PPI prediction algorithm based on CT features and Support Vector Machines (SVM), achieving an accuracy of 83.9%. With a consistent development of machine learning algorithms, the contribution of several other SVM [6][10][15-17], KNN [14], and random forest based methods were developed [18-21]. These extensive experiments have also indicated that appropriate feature extraction is one of the crucial steps in the path of the development of a robust predictor method. Another important step in that direction is the selection of an efficient dimension-reduction and noise-reduction technique. Most of the machine learning-based models have employed principal component analysis (PCA) that produces a discriminative feature subspace [22]. In recent years, an impressive set of methodologies based on deep neural networks have also been developed to predict PPIs [23-26] and nuclear receptors [27]. DeepPPI [25] and EnsDNN (Ensemble Deep Neural Networks) [28] are some methods that corroborate multi-descriptor techniques such as the combination of amino acid composition (AAC), conjoint triads (CT), auto-covariance descriptor (AC), Local descriptor and multi-scale continuous and discontinuous local descriptor (LD) & (MCD) to predict PPIs. The accuracy achieved by the DeepPPI and EnsDNN method, when executed over *Saccharomyces cerevisiae* datasets was 94.43%[25] and 95.29%[28] respectively. These methods achieved impressively higher accuracy than those of the traditional methods based on state-of-the-art machine learning algorithms but the execution of the model on the PPI networks was not elucidated in these studies. Since, the wet-lab experiments have identified only a fraction of complete PPI networks [29] therefore, a robust PPI predictory model is expected to predict the potential important PPI pathways and networks.

The inadequacies observed in the previous methods motivated us to develop a novel PPI prediction method based on the LightGBM classifier and Autoencoder, which is a powerful artificial neural network that is capable of reconstructing a large feature set into a smaller dimension of a discriminative and noise-free feature subspace. (a) In this paper, we firstly, constructed a combined feature set of the conjoint triad (CT) and Composition-Transition-Distribution (CTD) to reflect the complete structural and molecular properties of a protein into numeric feature vectors. (b) Secondly, an Autoencoder was employed to efficiently fuse the feature sets into a noise-free, lower-dimensional discriminative subspace. (c) Subsequently, a LightGBM, which is an open-source framework, was employed as a classifier. (d) Finally, the resultant prediction model was evaluated on other species PPI datasets and PPI pathways. **Fig 1** is a schematic representation of the AE-LGBM method. The configuration of the computer on which all experiments were carried out is Intel (R) Core(TM) i5-4310U, 8 GB RAM, and 64 bit OS Win 10. Method implementation and experiments are conducted using R 4.0.0 and Python 3.7.4.

**Fig 1:**
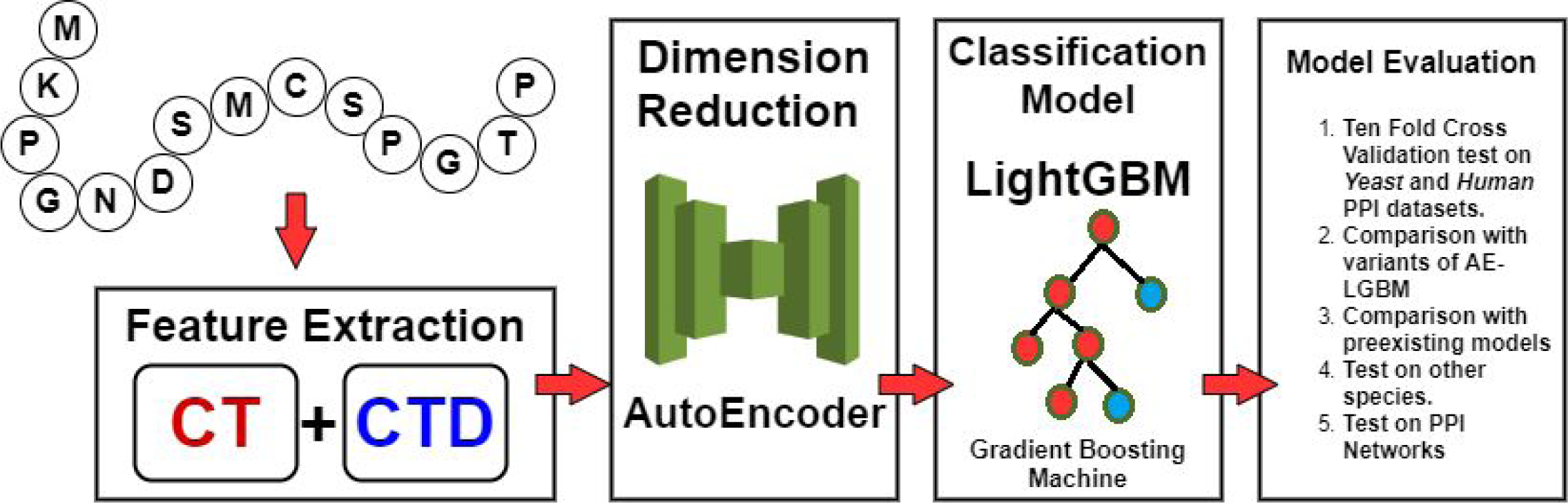
The complete framework of AE-LGBM

## 3. Methodology

### 3.1. Dataset

The method is constructed using the golden standard *Human* and *Saccharomyces cerevisiae (Yeast)* protein-protein interactions datasets, available on the opensource DIP database [30]. These datasets include 5,594 and 3,899 positive interacting protein pairs respectively. The negative interacting pairs were constructed using the previously defined standard strategies corroborated in the studies performed by Guo et al [31] and Shen et al [13]. Precisely, (1) Protein sequences of a shorter length (<50 amino acids) were eliminated from the pool. (2) Protein sequences with a pairwise sequence identity of more than 40% and 25% were removed for *Yeast* and *Human* respectively. (3) Assuming that the proteins belonging to different subcellular compartments like the nucleus, Golgi apparatus, cytoplasm, nucleus, etc are non-interacting, 5,594 non-interacting protein pairs are constructed for *Yeast*. In similar veins, a set of 4,262 negative PPI datasets is constructed for *Human* out of 661 positive interacting *Human* protein sequences fetched from Swiss-Prot (http://www.expasy.org/sprot/) [31]. This results in a total of 11,188 and 8,161, interacting and non-interacting instances of protein pairs for both Yeast and Human respectively. To evaluate the versatility of the AE-LGBM prediction model, we selected four standard test datasets namely, *Escherichia coli, C. elegan, Homo sapiens*, and *Mus musculus* containing 6,954, 4013, 1,412 and 313 PPI pairs respectively. To further measure the potential of our model, we implemented the same over three important PPIs networks namely (1) single-core network (CD9) containing 16 PPI pairs, (2) the multiple-core network (The Ras/Raf/MEK/ERK pathway) containing 198 PPI pairs, and (3) the cross-connection network (Wnt Network) containing 96 PPI pairs.

### 3.2. Feature Extraction

#### 3.2.1. Conjoint Triad

The efficiency of a PPI prediction model depends significantly on the competency of the features. The efficacy of a feature vector is its capability to express the entire sequence information into numerical vectors precisely. Conjoint Triad (CT) feature sets have demonstrated promising results in terms of mapping the interaction information of a protein sequence [13]. To extract a CT feature vector, 20 amino acids are classified into seven groups based on their dipole and side-chain volume (**Table S1**). Then sliding along the protein sequence, each amino acid is mapped to an integer, ranging from 1 to 7, based on their corresponding class number. For example, _***N***_MQNKLNKLLD_***C***_ is mapped as _***N***_7556456443_***C***_

To account for the interaction of amino acids with its adjacent amino acids in the protein sequence, the combination of three adjacent amino acids and their corresponding three-digit mapped numbers are considered as a triad. By sliding from *N*-terminal to *C*-terminal correspondingly on the mapped sequence with a window size of three residues and step size of one residue, the frequency (f_t_) of the t^th^ triad in the mapped sequence is computed, where t is a set of all unique triad i.e t= {111, 112, 113, …..776, 777}. The size of the set t is 7×7×7. For the given instance, we get

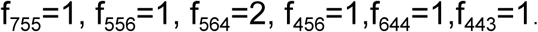

Finally, The CT-feature vector *CT* = (**f**_**111**_,**f**_**112**,_**f**_**113**_,**…..f**_**776**_,**f**_**777**_) of 343 dimensions is extracted from a single protein sequence (**Fig S1**).

#### 3.2.2. Composition-Transition-Distribution (CTD)

Much research in the past has indicated that the amino acids present in a protein sequence can be classified on the basis of structural and physicochemical properties like secondary structure and hydrophobicity [32-34]. Such structural and physicochemical properties of a protein sequence residue have been utilized to create advanced feature extraction toolkits [35]. We probe into one such multi-descriptor and employ it in our model. Composition-Transition-Distribution (CTD) is a multi-descriptor that includes a number of structural and physicochemical based attributes in which the residues of a protein sequence are grouped into three divisions (**Table S2**). To extract,

1. The *Composition*-feature vector, for each attribute, the amino acids are mapped into numbers ranging from 1 to 3, corresponding to their respective division(**Fig S2**). For instance, considering the attribute Hydrophobicity_FASG890101, the protein sequence **‘MQNKLNKLLD’** will then be represented as **‘3113221331’**, where ‘1’, ‘2’ and ‘3’ represents the ‘Polar’, ‘Neutral’ and ‘Hydrophobicity’ divisions respectively. Sliding along the protein chain from N terminal to C terminal, the relative frequency C_i_ of the i^th^ division is computed as f_i_/N, where f_i_ is the frequency of the i^th^ division, N is the total number of amino acids present in the protein sequence and i={1,2,3}. In this way, the *Composition* extracts a feature vector of 13×3 dimensions. For the given instance, C_1_=4/10, C_2_=2/10, C_3_=4/10.
2. The *Transition*-feature vector, the mapping strategy similar to *Composition* is employed on the sequence. We derive a transition matrix as:

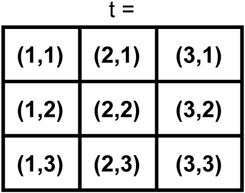

Sliding from the *N*-terminal to *C*-terminal with a window size of two residues and a step size of one residue, the frequency f of t_ij_ is calculated. The t_ij_ with i≠j were only considered as transitions and rest were ignored. Finally, *Transition*-feature vector T is extracted as:

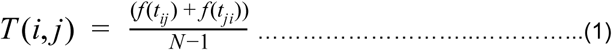

So,for the given instance, T(1,2) = 1/9, T(2,3)=1/9 and T(1,3)=4/9. A *Transition*-feature vector generated from all the attributes for a single protein sequence is of 13×3 dimensions.
3. The *Distribution*-feature vector, the position of the first, 25%, 50%, 75%, and 100% of each group is computed and divided by the total number of amino acids present in the protein sequence. This feature vector contains the information of the distribution pattern of the attribute-based different types of residues in the protein sequence. To extract the Distribution-feature for ‘3’ in ‘3113221331’, the position of first ‘3’ is 1. The position of 25% of ‘3’ is 0.25×4 =1. Likewise, the position of 50%, 75%, and 100% of ‘3’ is 4,8 and 9 respectively. For a single attribute, this process resulted in 3×5 features and similarly, the entire *Distribution*-feature vector of 195 dimensions was created for a single protein sequence.

### 3.3. AutoEncoder

The combination of CT and CTD feature vectors results in a high-dimensional (616) feature-set. Such a high-dimensionality can lead to redundancy, computational complexity, and overfitting [36]. To overcome this issue, we employed an Autoencoder which is a generative neural network. Autoencoders are a capitative dimension reduction tool that has a great potential to handle noisy datasets. Deep learning algorithms have undergone tremendous advancements over the decades [37]. Autoencoders are now coming up as powerful alternatives for standard dimension reduction techniques like PCA and have contributed to breakthrough discoveries in computational biology [24] [38-40]. .An Autoencoder is a generative neural network that is made up of an encoder and a decoder which together are joined through a hidden layer or latent of a smaller dimension [41]. The encoder assimilates the high-dimensional input data and passes it through the latent to the decoder for a reconstruction (**Fig 2**). Precisely, the underlying mathematical functions(eq 2-3) and optimization objective (eq 4) behind the encoder and decoder of an Autoencoder are:

**Fig 2.**
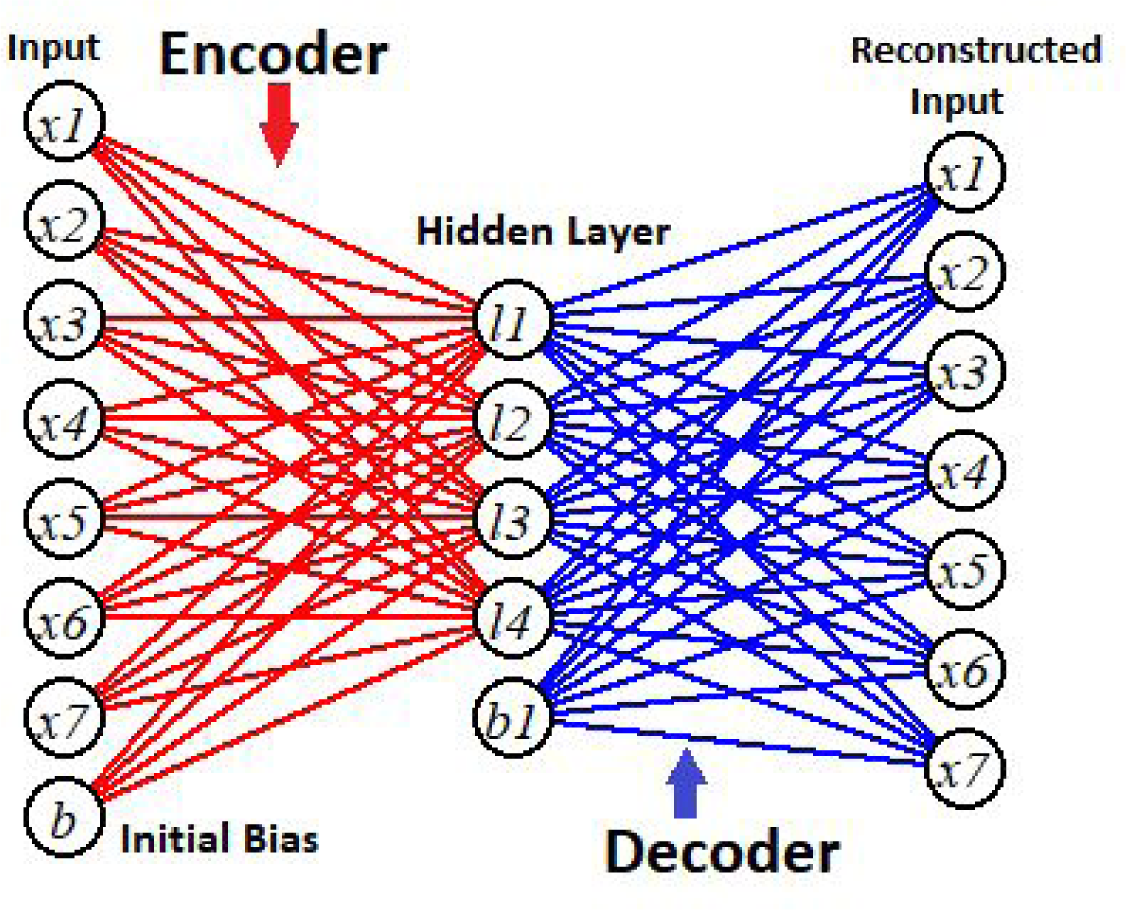
An Autoencoder with one hidden layer

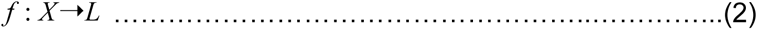

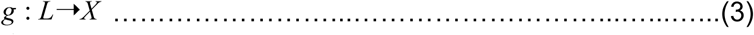

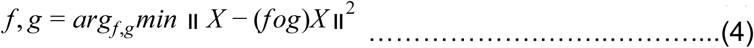

Where, *f* is an encoder that transforms the input *X* into the latent layer, and subsequently, the decoder function *g*, morphs the latent space *L* into the output in order to reconstruct the input following the optimization goal (eq 4).

Training of an Autoencoder is done by initializing the weight (*W, W* ′) and bias (*b, b*′) for both the encoder and decoder respectively. The scheme then follows:

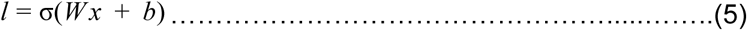

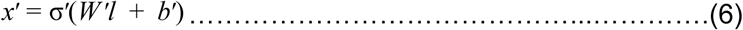

Where σ and σ’ are activation functions, we employ sigmoid functions as activation functions. The loss function ɀ is then defined as:

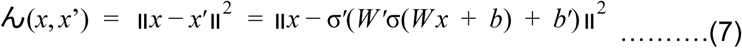

The latent transforms the high-dimensional feature vector into a discriminative and noise-free lower condensed feature space. An appropriately fine-tuned autoencoder through gradient descent can perform superior to principal component analysis (PCA) [41] (**Fig S3**) The complete description of the configuration of our Autoencoder is provided in addition material (**Table S3**)

### 3.4. LightGBM

J.H. Friedman [42] demonstrated that a gradient boosting algorithm attains a highly competitive and robust accuracy for both classification and regression. Based on it, Chen et al [43] proposed a scalable tree boosting machine called XGBoost. Although XGBoost had attained excellent accuracy, the LightGBM ensemble has achieved the same but has been more robust, less computationally expensive, and time-efficient [44]. A brief and precise explanation of the gradient boosting decision tree (GBDT) algorithm is found in Cheng et al [45]. For a given training set *X* = {(*x*_*i*_, *y*_*i*_) | *x*_*i*_ ∈ *R*^*k*^, *y*_*i*_ ∈ *R, k*≥1, |*X*| = *n*} where x is the features and y is the label, considering *F*_*o*_ as an initial fit and ɀ be the loss function, the optimization goal (eq 8) is given as,

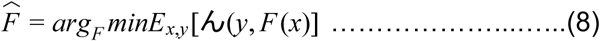

The *pseudo* residuals or gradient α_im_ for the m^th^ iteration is obtained as (eq 9), to which the decision tree *h*_*m*_(*x*) is fitted,

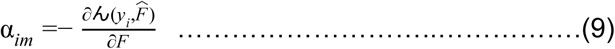

In order to minimize the loss function, the resultant iterative criterion for GBDT to obtain a new boosted fit is (eq 10),

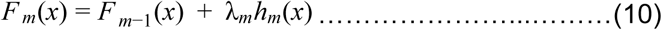

Where 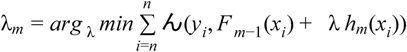 is a multiplier that serves as step size and is optimized through a linear search. In this paper, we employ the LightGBM tool to construct the PPI prediction model. LightGBM is a robust gradient boosting framework with a base classifier as a decision tree. The description of the best performing parameters in our case can be found in the additional material (**Table S4**). LightGBM package is available at *https://github.com/Microsoft/LightGBM*.

### 3.5. Performance evaluation

To evaluate the predictive performance of our ensemble model, we performed 10-fold cross-validation, considering that complex gradient boosting models are prone to overfit on a small training sample [46]. The randomly shuffled dataset was divided into ten equal sets and the evaluation was conducted by testing the model on a set in turn, while trained over the remaining of the nine sets. The average performance over the ten sets was considered as the performance of the AE-LGBM. We use four standard evaluation indicators to measure the model’s predictive potentials. These indicators are prediction accuracy (Acc), sensitivity (Sn), specificity (Sp), and Matthew’s correlation coefficient (Mcc). We computed these metrics using the standard definitions [22] which are as follows:

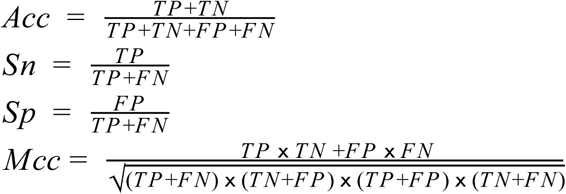

Where TP, TN, FP, and FN stand for true positive, true negative, false positive, and false negative respectively. The receiver operating characteristic curve (ROC) along with its area under the curve (AUC) is also elucidated to compare our model with its variants and other previous state-of-the-art prediction models. A ROC curve is a plot of true positive rate against the false-positive rate under a varying threshold. The plot elucidates a tradeoff between the sensitivity and specificity of the model and the area under the ROC curve, AUC indicates the usefulness of the model where a larger area indicates a higher predictive ability. ROC curves containing a sharp bending towards the upper-left corner reflects a better performance, indicating that the model achieves high sensitivity with a lower false-positive rate. Cross-species predictive potential of AE-LGBM was evaluated on other species PPI datasets and PPI pathways while assimilating the *Human* PPI dataset as the training set.

## 4. Results and Discussion

The AE-LGBM ensemble invites a good deal of hyperparameter tuning. To address this, we focused on the most sensitive hyperparameters. The parameters evincing a negligible impact on performance in the first few experiments were kept static. Description of parameters are provided in additional material (**Table S3-S5**). An autoencoder with a single hidden layer was sufficient to attain the best prediction accuracy of 96.7% and 99.4% for both Yeast and Human PPI data sets respectively at a fairly lower computation cost. Interestingly, such phenomena have been reported in previous studies as well, where adding more than one layer to an autoencoder was futile [24,38]. For the Auto-Encoder and LightGBM, the number of nodes in the latent (N) and the maximum number of leaves (Max_L) were fine-tuned. We evaluated the performance of AE-LGBM for different combinations of N and Max_L. N varied in the range of 200 to 209 and the Max_L was set to 50, 80, 110, 150, and 200 in turn for each N. Comprehensive graphical representations of tuning results are elucidated (**Fig 3)**. The best 10 fold-CV prediction accuracy for Human PPIs (98.65%) was achieved at setting the N to 208 and Max_L to 80 (Fig.3 A, B). For the Yeast dataset, the best accuracy (95.37%) was attained at 202 nodes and at maximum leaves of 50 (Fig.3 C, D). As expected, these configurations have attained the highest sensitivity, specificity, and Mcc among all the other combinations. Comparative bar graphs of results obtained at different nodes and max_leaf are added in the additional material (**Fig S4**). The average AUC for all ten-folds cross-validation were 0.998 and 0.987 for both Yeast and Human respectively, which is higher than other state-of-the-art models like EnsDNN (0.975 for Yeast) [23], DNNs+APAAC (0.975 for Yeast) [25] and WSRC+PSM (0.971 for Yeast) [47] (**Fig4**).

**Fig 3.**
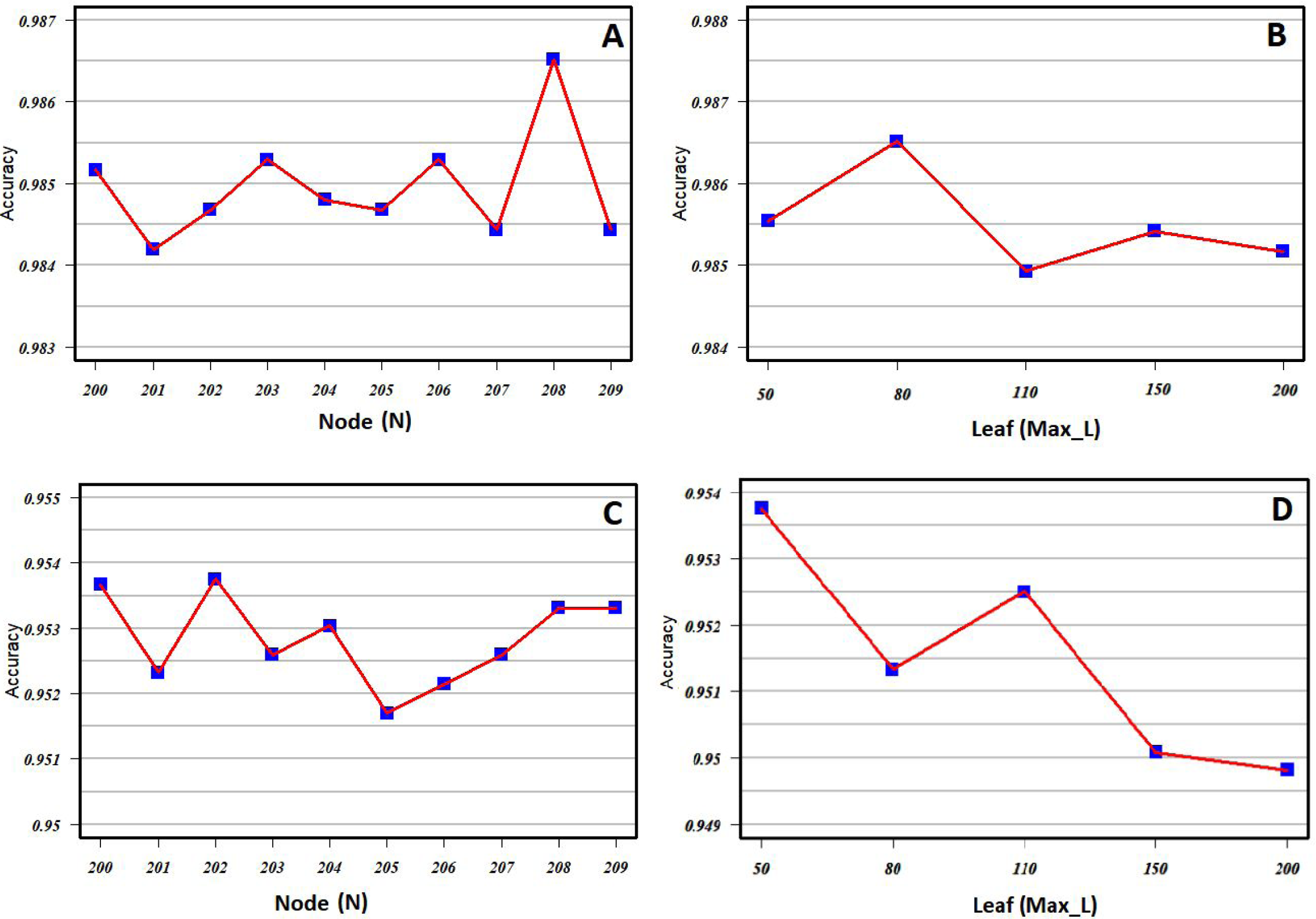
The accuracy obtained by AE-LGBM at different numbers of nodes and maximum leaf. (A) and (B) represent the accuracy obtained for Human PPIs by AE-LGBM at varying numbers of nodes (N) and maximum leaves (Max_L). The maximum accuracy was observed at 208 N and 80 Max_L. (C) and (D) represent the accuracy obtained for YeastPPIs by AE-LGBM at varying numbers of nodes (N) and maximum leaves (Max_L). The maximum accuracy was observed at 202 N and 50 Max_L.

**Fig 4.**
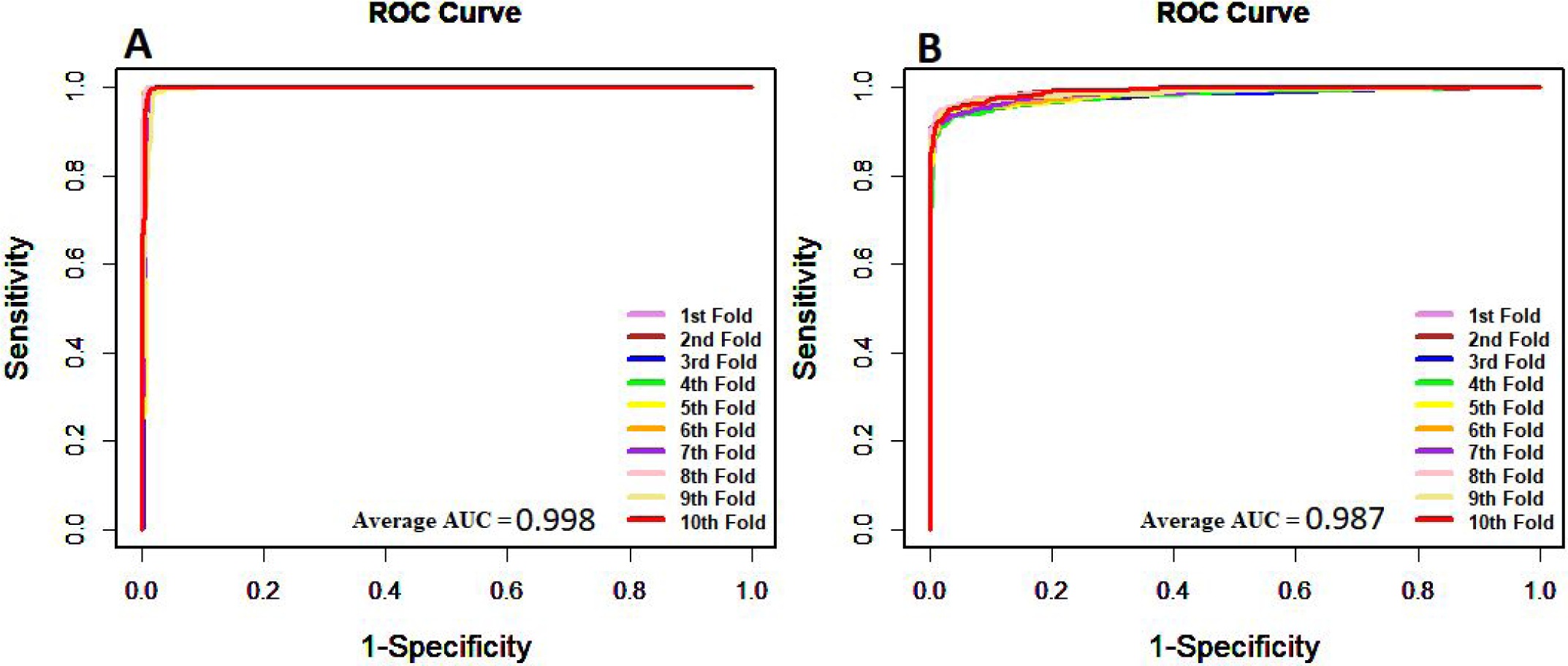
**(A)** ROC curve of AE-LGBM on Human PPI dataset. **(B)** ROC curve of AE-LGBM on Yeast PPI dataset.

To further evaluate the competency of AE-LGBM, we compared the model’s performance with some other state-of-the-art classifiers namely support vector machine (SVM) and random forest (RF). We built two ensembles, AE-SVM and AE-RF, by configuring a single-layered Autoencoder and state-of-the-art classifiers for both Human and Yeast PPIs. Descriptions of the customized parameters of SVM and RandomForest are added in additional material (**Table S5**). In these ensembled models, the Autoencoder contained the optimal number of nodes N in their hidden layer, that were found for Yeast(202) and Human(208) cases. We then performed a 10-fold-CV with these models on the PPI datasets. The AE-LGBM significantly outperformed AE-SVM and AE-RF(**Fig5**). The sensitivity and the precision obtained by AE-LGBM were also significantly higher than that of AE-SVM and AE-RF models (**Table 1**). The AUC achieved by AE-LGBM (0.998 for Human and 0.987 for Yeast) were higher than the AUC obtained by AE-SVM (0.997 for Human and 0.982 for Yeast) and AE-RF (0.998 for Human and 0.979 for Yeast) (**Fig6**).

**Table 1.** Performance comparison of AE-LGBM with AE-SVM and AE-RF for both Human and Yeast PPI datasets

| Dataset | Model | Accuracy | Specificity | Sensitivity | Mcc |
| --- | --- | --- | --- | --- | --- |
| *Human* | **AE-LGBM** | **0.987±0.001** | **0.992±0.002** | **0.981±0.002** | **0.973±0.003** |
|  | AE-SVM | 0.973±0.002 | 0.985±0.002 | 0.959±0.003 | 0.946±0.004 |
|  | AE-RF | 0.985±0.002 | 0.99±0.002 | 0.98±0.003 | 0.97±0.004 |
| *Yeast* | **AE-LGBM** | **0.954±0.002** | **0.987±0.002** | **0.921±0.003** | **0.91±0.004** |
|  | AE-SVM | 0.934±0.002 | 0.963±0.003 | 0.906±0.004 | 0.87±0.004 |
|  | AE-RF | 0.943±0.002 | 0.977±0.002 | 0.908±0.004 | 0.887±0.004 |

**Fig 5.**
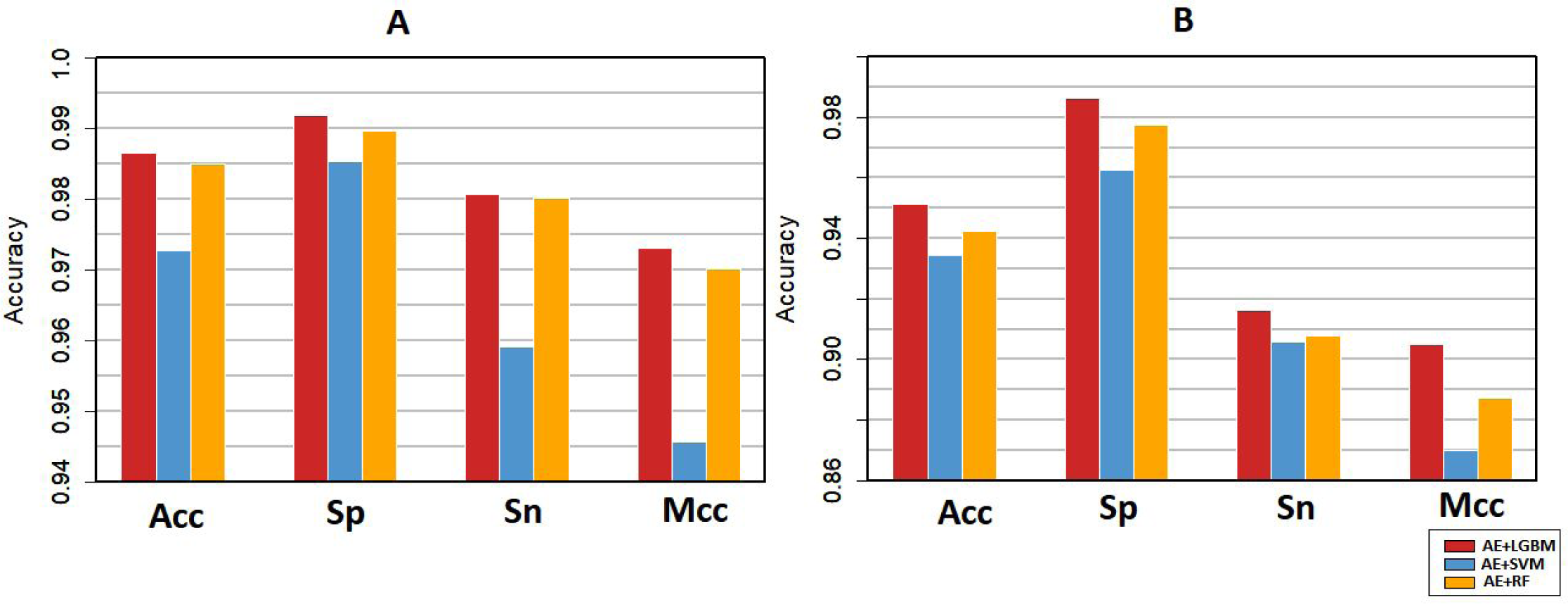
**(A)** Represents the performance indicators of AE-LGBM, AE-SVM and AE-RF on Human PPI dataset. **(B)** Represents the performance indicators of AE-LGBM, AE-SVM and AE-RF on Yeast PPI dataset. Acc is accuracy, Sp is specificity, Sn is sensitivity and Mcc is Matthew’s correlation coefficient.

**Fig 6.**
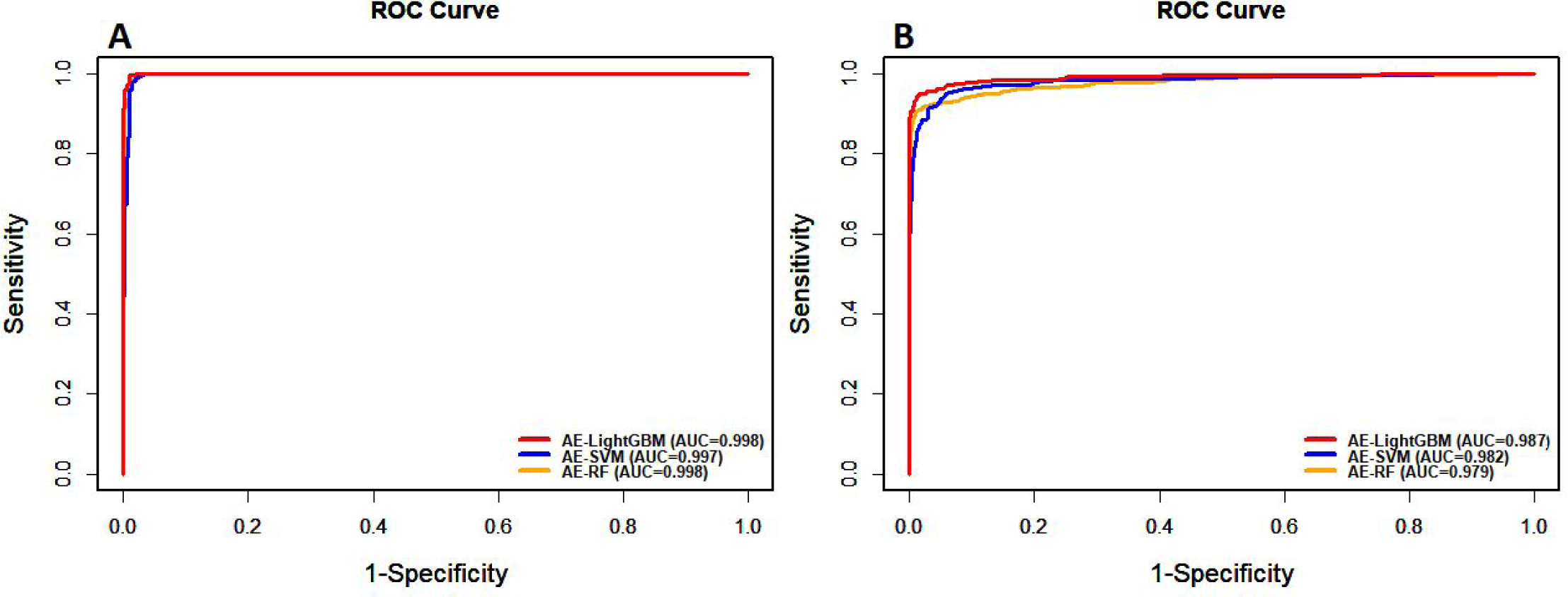
**(A)** ROC curve of AE-LGBM, AE-SVM, and AE-RF on Human PPI dataset. **(B)** ROC curve of AE-LGBM, AE-SVM, and AE-RF on Yeast PPI dataset.

We also compared the predictive abilities of our model with some preexisting remarkable PPI prediction models. **Table. 2** indicates that the accuracy attained by AE-LGBM on Human PPIs (98.65%) was 1.01% higher than WSRC+GE [48], 1.46% higher than AE-AC[24], 2.25% and 3.0% higher than LDA+RF [49] and LDA+RoF[48]^]^ respectively. AE-LGBM has outperformed autocorrelation (AE), based models, by scoring 3.1% and 3.5% higher accuracy than that of AC+RF and AC+RoF models [49]. **Table. 3** accounts the performance indicators of eight other prediction models on Yeast dataset including EN-LGBM [44], DeepPPI [25], HOG+SVD+RF [50], RVM+BiGP [51], RF+PR+LPQ [52], AutoCC [31], SVM+LD [53] and PCA+ELM [22].

**Table 2.** Performance comparison of AE-LGBM with other PPI prediction models on Human PPIs dataset. N/A means the value of the indicator is not queried.

| Dataset | Model | Accuracy | Sensitivity | Specificity | Mcc |
| --- | --- | --- | --- | --- | --- |
| *Human* | **AE+LGBM** | **0.987** | **0.981** | **0.992** | **0.973** |
|  | WSRC+GE | 0.9766 | 0.9528 | 0.9981 | 0.9541 |
|  | AE-AC | 0.9719 | 0.9806 | 0.9634 | N/A |
|  | LDA+RF | 0.964 | 0.942 | N/A | 0.928 |
|  | LDA+RoF | 0.957 | 0.976 | N/A | 0.918 |
|  | AC+RF | 0.955 | 0.94 | N/A | 0.914 |
|  | AC+RoF | 0.951 | 0.933 | N/A | 0.91 |

**Table 3.** Performance comparison of AE-LGBM with other PPI prediction models on Yeast PPIs dataset. N/A means the value of the indicator is not queried.

| Dataset | Model | Accuracy | Sensitivity | Specificity | Mcc |
| --- | --- | --- | --- | --- | --- |
| *Yeast* | **AE+LGBM** | **0.954** | **0.987** | **0.921** | **0.91** |
|  | EN+LGBM | 0.9507 | 0.979 | 0.922 | 0.903 |
|  | DeepPPI | 0.95 | 0.944 | 0.92 | 92.21 |
|  | HOG+SVD+RF | 0.9483 | 0.924 | 0.971 | 0.8977 |
|  | RVM+BiGP | 0.9457 | 0.9427 | 0.9486 | 0.8974 |
|  | RF+PR+LPQ | 0.9392 | 0.911 | 0.9645 | 0.8856 |
|  | AutoCC | 0.8933 | 0.8993 | 0.8887 | N/A |
|  | SVM+LD | 0.8856 | 0.8737 | 0.895 | 0.7715 |
|  | PCA+ELM | 0.87 | 0.8615 | 0.8759 | 0.7736 |

In order to evaluate the generalizability of AE-LGBM, we trained the AE-LGBM on the entire Human data set and tested it on *E.coli, M.musculus, C.elegans*, and *H.sapiens* PPIs. AE-LGBM achieved an impressive accuracy of 100%,100%, 99.9%, 99.2% for *E.coli, M.musculus, C.elegans*, and *H.sapiens* respectively. To our knowledge, this is the highest prediction cross-species accuracy achieved by any preexisting PPI model (**Table 4**). This shows that AE-LGBM is a powerful cross-species PPI prediction model.

**Table 4.** Accuracy comparison of AE-LGBM with other prediction models on cross-species PPIs datasets. EN+LGBM [44], DeepPPI [25], DCT WSRC[54] and SVM [55].

| Species | AE+LGBM |  |  | EN+LGBM | DeepPPI | DCT WSRC | SVM |
| --- | --- | --- | --- | --- | --- | --- | --- |
|  | TP | FN | Acc | Acc | Acc | Acc | Acc |
| *E.coli* | 6953 | 0 | **1.00** | 0.921 | 0.921 | 0.66 | 0.712 |
| *M.musculus* | 313 | 0 | **1.00** | 0.945 | 0.913 | 0.798 | 0.766 |
| *C.elegans* | 4012 | 1 | **0.999** | 0.901 | 0.948 | 0.811 | 0.757 |
| *H.sapiens* | 1402 | 10 | **0.992** | 0.948 | 0.937 | 0.822 | 0.762 |

A good prediction ability of PPI networks should be present in the toolbox of every robust PPI prediction model. Although, a number of models were created to predict the PPI networks but remained unsatisfactory. We tested AE-LGBM abilities on three important PPI networks including (1) single-core network (CD9) consist of 16 genes interacting with a core protein CD9, (2) The multiple-core network (The Ras/Raf/MEK/ERK pathway) consists of 198 readily interacting protein pairs and (3) based on Wnt related pathway, the cross-connection network of 96 interacting protein pairs.

CD9 is a tetrameric protein ensemble that is widely distributed across various human body tissues. CD9 plays a vital role in tumor suppression, cell motility, adhesion, and other cellular fusions including sperm-egg fusion [56]. The AE-LGBM was able to identify all 16 PPIs involved in the CD9 network, giving an impressive accuracy of 100% (**Fig 7 A**). The complex mitogen-activated protein (MAP) kinase pathway activates the Ras-Raf-Mek-Erk-Elk-Srf pathway which is responsible for signal transduction to the nucleus through growth factor receptors. Mostly found in eukaryotic cells, the Ras-Raf-Mek-Erk-Elk-Srf pathway is closely associated with tumor growth [57]. All the 196 protein pairs involved in this pathway were accurately identified by AE-LGBM with an outstanding performance of 100% accuracy (**Fig 7 B**). A majority of PPIs networks are cross-connection networks. A robust prediction model is an unmet requirement for such networks. We tested AE-LGBM on the Wnt signaling pathway that is composed of multi-core networks and is closely related to tumor growth [58]. The AE-LGBM was able to predict 97 out of 98 interacting pairs with remarkable accuracy of 98.9% (**Fig 7 C**). To our knowledge, the overall performance of AE-LGBM over these three PPI networks is superior to any existing PPI prediction model.

**Fig 7(A).**
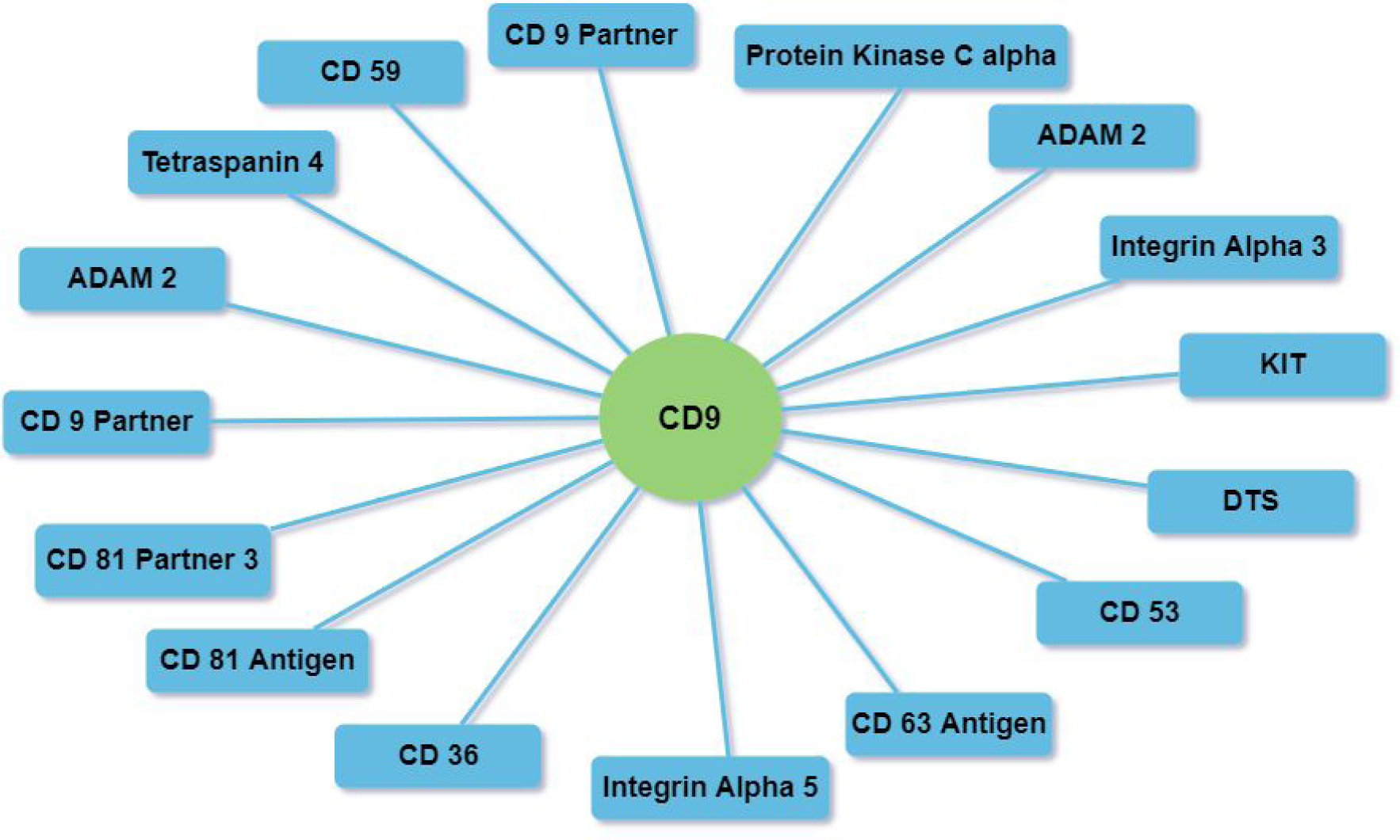
Prediction result of AE-LGBM on Single core CD9 network

**Fig 7(B).**
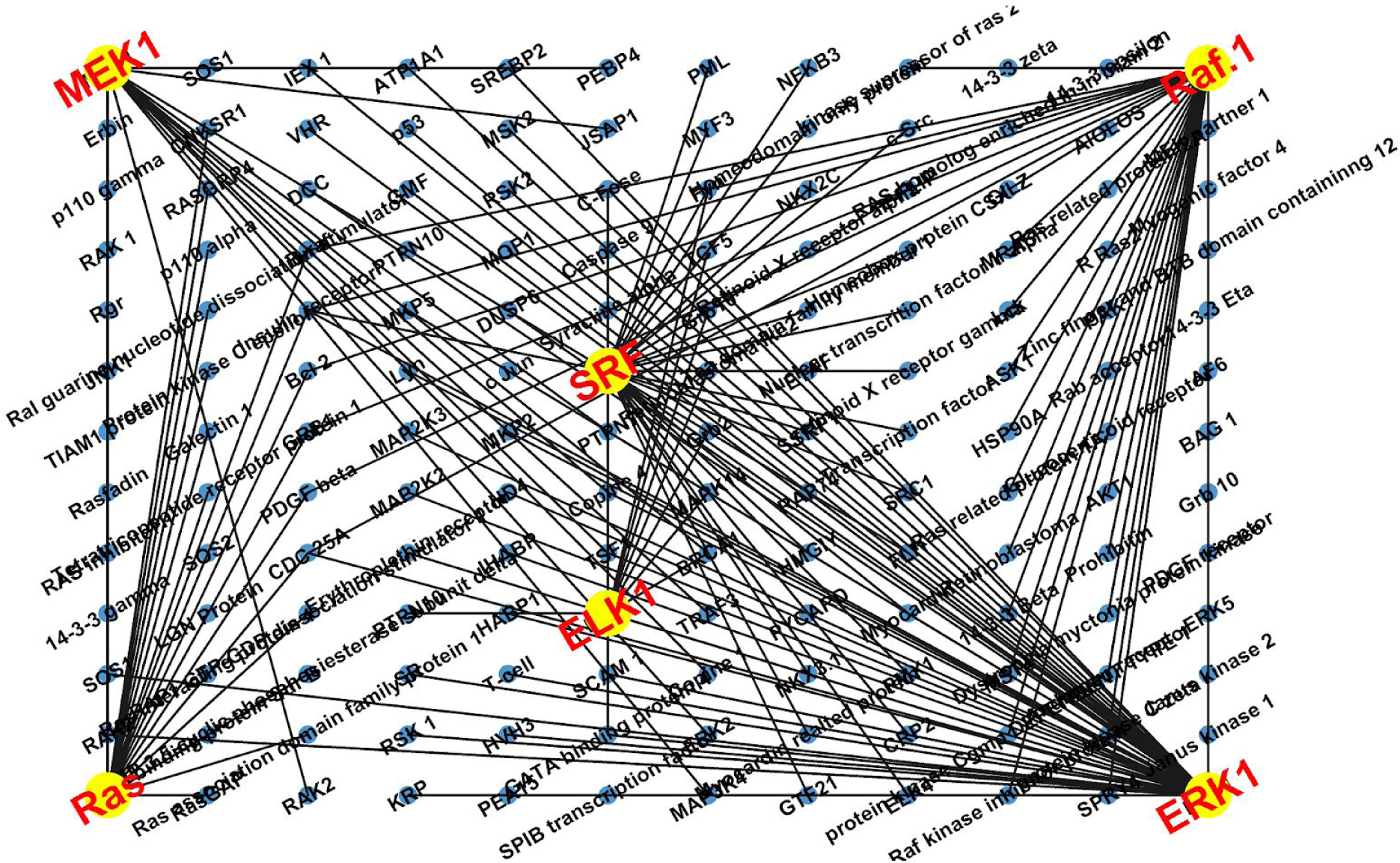
Prediction result of AE-LGBM on Ras-Raf-Mek-Erk-Elk-Srf pathway

**Fig 7(C).**
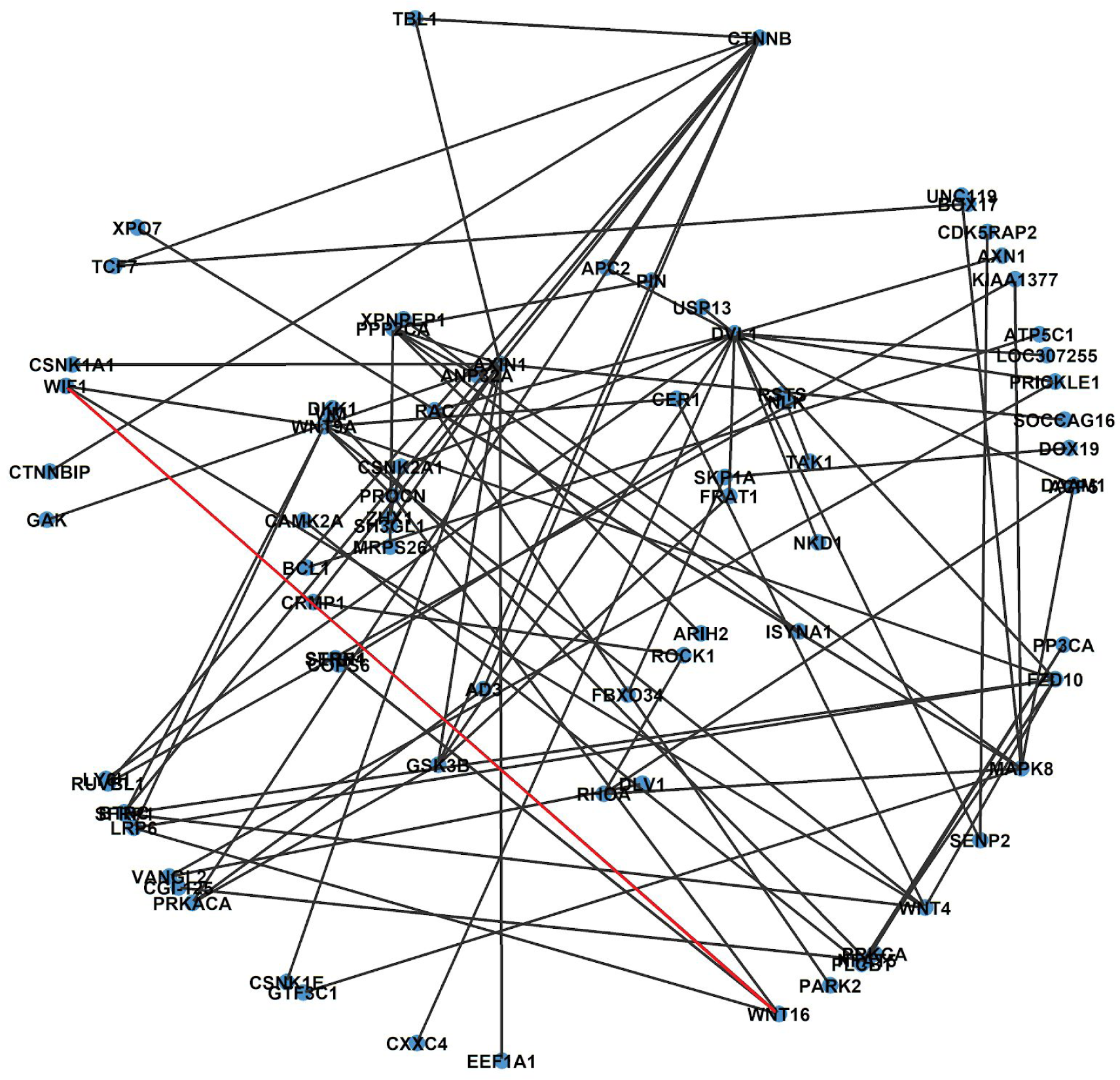
Prediction result of AE-LGBM on Wnt - Network. Red line indicates the false prediction.

## 5. Conclusion

In biology, The prediction of protein-protein interactions (PPIs) is identified as a key mechanism behind every biological process. A computer-based prediction model to predict PPI has important practical significance. In this paper, we proposed an Autoencoder and LightGBM based predictory model AE-LGBM, that tackles the prediction objective robustly and intensively at a fairly lower cost of time and computation, giving the best overall performance than any other preexisting machine learning based prediction model. We utilized the sequence information as features, which was extracted using conjoint triad (CT) and composition, transition, and distribution (CTD) feature extraction methods and condensed to a lower dimension using Autoencoder. When trained and tested on the gold standard S. cerevisiae and Human PPI dataset, the model achieved excellent accuracy. AE-LGBM has also outperformed its own variants including SVM based AE-SVM and random-forest based AE-RF. AE-LGBM attained so far the best accuracy on cross-species PPI datasets including E.coli, M.musculus, C.elegans, and H.sapiens. We evaluated AE-LGBM on three important PPI networks CD9, The Ras/Raf/MEK/ERK pathway, and Wnt-network. AE-LGBM predicted all the interacting pairs except one pair, which is a highly competitive performance.

AE-LGBM is a versatile PPI prediction model, which can potentially serve as an important tool to discover new insights into vital biological processes. AE-LGBM amalgamate neural network architecture with a revolutionary decision tree as a base classifier. This opens a new window for enhancement in performance as every year, new insights are being developed in the formalism of these tools. To bring AE-LGBM into the public domain for the benefit of the scientific community, our next step would be to compile AE-LGBM into a robust package and distribute it on various computing platforms like R and Python.

